# Tuning the immunostimulation properties of cationic lipid nanocarriers for nucleic acid delivery

**DOI:** 10.1101/2021.06.16.448666

**Authors:** Arindam K Dey, Adrien Nougarede, Flora Clément, Carole Fournier, Evelyne Jouvin-Marche, Marie Escudé, Dorothée Jary, Fabrice P. Navarro, Patrice N Marche

## Abstract

Nonviral systems, such as lipid nanoparticles, have emerged as reliable methods to enable nucleic acid intracellular delivery. The use of cationic lipids in various formulations of lipid nanoparticles enables the formation of complexes with nucleic acid cargo and facilitates their uptake by target cells. However, due to their small size and highly charged nature, these nanocarrier systems can interact *in vivo* with antigen-presenting cells (APCs), such as dendritic cells (DCs) and macrophages. As this might prove to be a safety concern for developing therapies based on lipid nanocarriers, we sought to understand how they could affect the physiology of APCs. In the present study, we investigate the cellular and metabolic response of primary macrophages or DCs exposed to the neutral or cationic variant of the same lipid nanoparticle formulation. We demonstrate that macrophages are the cells affected most significantly and that the cationic nanocarrier has a substantial impact on their physiology, depending on the positive surface charge. Our study provides a first model explaining the impact of charged lipid materials on immune cells and demonstrates that the primary adverse effects observed can be prevented by fine-tuning the load of nucleic acid cargo. Finally, we bring rationale to calibrate the nucleic acid load of cationic lipid nanocarriers depending on whether immunostimulation is desirable with the intended therapeutic application, for instance, gene delivery or messenger RNA vaccines.

## 1 Introduction

In recent years, advances in field of nanotechnology have demonstrated potential for precision medicine. For instance, lipid nanoparticles (LNPs) can be used for the targeted delivery of therapeutic molecules, increasing their bioavailability and pharmacokinetic properties beyond the Lipinski rules (Yousefi and Tufenkji, 2016). Indeed, the development of nucleic acid therapeutics has long been hampered by the inherent hydrophilic nature, large size, and poor membrane permeability of nucleic acids (Stoddard et al., 2018). LNPs can be a potent alternative to viral-mediated nucleic acid delivery, with an extensive range of applications such as RNA interference (RNAi) therapy or RNA-based vaccines through intracellular delivery, respectively, of short interfering RNA (siRNA) or messenger RNA (mRNA) (Xue et al., 2015).

One of the primary advantages associated with LNPs is their biocompatibility that enables their use *in vivo* for human therapy (Chira et al., 2015;Hu et al., 2020). LNPs are made of two major components: a lipid phase and a water phase containing surfactants. LNPs are generally divided into liposomes with an aqueous core or other LNPs; the latter could be solid lipid nanoparticles (SLNs) with a solid core and nanostructured lipid carriers (NLCs) featuring a core that is a mixture of solid and molten lipids (Mehnert and Mäder, 2001). This subclass of LNPs was initially designed to improve the colloidal stability of lipid carriers and increase the drug payload into the core by controlling the release profile (zur Mühlen et al., 1998). Moreover, they are considered advantageous because their manufacturing processes can be easily scaled up for large production (Müller et al., 2002).

Due to the nature of their lipid core, these particles are not well adapted for nucleic acid encapsulation. The loading of biomacromolecules such as siRNA or mRNA, therefore, occurs through the association with their shell either by chemical modifications of Polyethylene glycol (PEG) residues (Kim et al., 2008) or by incorporation of cationic lipids at the level of phospholipid monolayer, thus allowing electrostatic interactions with negatively charged nucleic acids (del Pozo-Rodríguez et al., 2007;Kim et al., 2008;Taratula et al., 2013;Resnier et al., 2014). The most chosen cationic lipids are quaternised cationic lipids, such as Dioleoyl-3-trimethylammonium propane (DOTAP), which are added to the formulation at the appropriate ratio (Bruniaux et al., 2014). The NLCs with DOTAP present thereby a globally positive charge; thus, their toxicity and their impact on the immune systems need to be assessed. A previous study has reported that positively charged nanocarriers induce some systemic toxicity and pro-inflammatory effects (Kedmi et al., 2010). The microenvironment is known to drive distinct antigen-presenting cell (APC) fates by affecting functions of macrophages and dendritic cells (DCs) by activating different metabolic pathways. For example, while lipopolysaccharides (LPS) classically activated macrophages (M1), displaying pro-inflammatory activity, rely on glycolysis, Interleukin 4 (IL-4) alternatively activated macrophages (M2), displaying anti-inflammatory activity, primarily utilise fatty acid oxidation (FAO) and oxidative phosphorylation (OXPHOS) (Stunault et al., 2018). DCs, like macrophages, respond differently in the presence of LPS and IL4 (Wculek et al., 2019).

The exposition to cationic lipid carriers (cNLCs) has been shown to affect the functions of APCs. For instance, cNLCs were shown to activate bone-marrow-derived dendritic cells (BMDCs) partially by inducing the expression of two costimulatory molecules, CD80 and CD86, but without inducing the secretion of pro-inflammatory cytokines (Vangasseri et al., 2006).

DOTAP itself could interact directly with ligands on the surface of the immune system (de Groot et al., 2018). In the cationic NLCs formulation, we describe here that the phospholipid layer incorporating cationic lipids is covered by a dense PEGylated coating that contributes to the stability and also is known to reduce the interaction with proteins and other biological entities (Nel et al., 2009;Kedmi et al., 2010;Blanco et al., 2015).

Moreover, how the positive charge of lipid particles modulates the metabolic fitness of APCs and how this is related to the cellular function have not yet been elucidated. Therefore, understanding the impact of positively charged particles on immune responses and particularly on APCs metabolism, fate and cytokine secretion is crucial to control the use of nanocarriers fully.

In the present study, we analysed the effect of NLCs surface charge on primary APCs using BMDCs and bone-marrow-derived macrophages (BMDMs), as cellular models. We evaluated the impact of neutral lipid carriers (nNLCs) and cNLCs on the secretion of different signalling factors and mitochondrial metabolism and glycolysis. Furthermore, we used negatively charged siRNA to reverse the net charge on cNLCs and evaluate the effect of different surface charges on cell function.

## 2 Materials and Methods

### 2.1.1 Cell culture

The murine macrophage cell line (J774.1A) was purchased from ATCC; the cells were cultured in Dulbecco’s modified Eagle’s medium (DMEM) supplemented with 10% fetal bovine serum and 1% penicillin-streptomycin.

As previously described (Faure et al., 2004), BMDCs were generated from the bone marrow extracted from C57BL/6 mice (Charles River, l’Arbresle, France). Bone marrow cells were isolated by flushing from the tibia and femur. Erythrocytes and GR1 positives cells were removed by incubating with Ly-6G/Ly-6C (BD Pharmingen, #553125) and TER-119 (BD Pharmingen, #553672) antibodies, and the remaining negatively sorted cells were isolated using Dynabeads isolation kit (ThermoFisher, #11047) by magnetic cell sorting; then the remaining negatively sorted cells were resuspended at 5×10^5^ cells/ml in complete Iscove’s modified Dulbecco’ s medium supplemented with Granulocyte-macrophage colony-stimulating factor (GM-CSF) (PeproTech, #315-03), FLT-3L (PeproTech, #250-31L) and Interleukin 6 (IL-6) (Peprotech, #216-16) according to Table 1. The transformation of the progenitors into fully active DCs was performed over a 10-day time frame.

**Table 1:** Concentration of GM-CSF, FLT-3L and IL-6 for BMDCs culture. Culture of BMDCs: BMDCs were seeded into a 100-mm TC-treated cell culture dish with 15 mL culture media. Culture media is supplemented with variable concentrations of GM-CSF, FLT-3L and IL-6 on day 0, day 3, day 5, day 7 and day 10 to harvest fully differentiated BMDCs on day 11.

| Cells are cultured in a 100-mm TC-treated cell culture dish with 15 mL culture media |  |  |  |  |  |  |
| --- | --- | --- | --- | --- | --- | --- |
|  |  | Day 0 | Day 3 | Day 5 | Day 7 | Day 10 |
| Cell concentration | | $0.6 \times 10^6/\text{mL}$ | $0.5 \times 10^6/\text{mL}$ | $0.5 \times 10^6/\text{mL}$ | $0.5 \times 10^6/\text{mL}$ | According to cell plating |
| Supplement | IL-6 | 5 ng/mL | 2.5 ng/mL | 2.5 ng/mL | – | – |
|  | FLT-3L | 50 ng/mL | 40 ng/mL | 30 ng/mL | 25 ng/mL | 25 ng/mL |
|  | GM-CSF | 5 ng/mL |  |  |  |  |

BMDMs were also generated from bone marrow extracted from C57BL/6 mice as previously described (Chen et al., 2021). Briefly, the erythrocytes were removed by the RBC lysis buffer, and the remaining cells were cultured in a complete DMEM with 20% L929 (Sigma, #85011425) in conditioned medium (source of macrophage colony-stimulating factor) for 7 days.

### 2.1.2 Cationic and neutral lipid nanocarriers

nNLCs and cNLCs were prepared as described in the previous study (Courant et al., 2017). Briefly, for nNLCs, a lipid phase was prepared containing triglycerides (Suppocire NB, Gattefossé and super-refined soybean oil, Croda Uniqema) and phospholipids (Lipoid SPC3, Lipoid). For cNLCs, the same lipid phase supplemented with the cationic lipid DOTAP (1,2-dioleoyl-3-trimethylammonium-propane chloride, Avanti Polar Lipids) and fusogenic lipid DOPE (1,2-dioleoyl-sn-glycero-3-phosphoethanolamine, Avanti Polar Lipids) were used.

When indicated, Dil lipophilic dye (D282, ThermoFisher) was added to the lipid phase to enable fluorescence detection of nNLCs. A second aqueous phase containing the PEGylated surfactant PEG-40 Stearate (Myrj S40, Croda Uniqema) was prepared in Phosphate-buffered saline (PBS) (#806552, Sigma). Both lipid and aqueous phases were mixed together through high-frequency sonication. Lipid nanoparticles are purified by dialysing in 100 volumes of LNP buffer: 154 mM NaCl, 10 mM HEPES, and pH 7.4 using endotoxin-free ultra-pure water (TMS-011-A, Sigma) and 12–14 kDa MW cut-off membranes (ZelluTrans/Roth T3). Finally, the LNP solution was sterilised by filtrating through a 0.22-μm millipore membrane.

### 2.1.3 Nanoparticle uptake assay

For nanoparticle uptake assays, 0.5 x 10^5^ cells/mL of BMDCs and BMDMs were seeded into a 4-well Lab-Tek chambered coverslip. After 24 h of growth, the cells were incubated with both Dil-labelled nanocarriers, cNLCs and nNLCs, for 24 h at 37°C with 5% CO_2_. Nanocarrier accumulation inside cells was monitored by time-lapse microscopy using a spinning disk confocal microscope (Andromeda, TILL-FEI). The Dil-labelled nanocarriers were visualised using the lipophilic dye excitation wavelength of 514 nm while plasma membranes were labelled with FITC-conjugated cholera toxin (Sigma, C1655) and visualised at the excitation wavelength of 488 nm. After acquisition, the images were processed in Icy 2.0.3.0 software, and spectral deconvolution was performed using NIS 5.20.01 software.

### 2.1.4 Physical characterisation of NLCs

The hydrodynamic diameter and polydispersity index (PDI) of the NLCs were determined by dynamic light scattering (DLS), and the zeta potential was determined by electrophoretic light scattering (ELS) using a Zetasizer Nano ZS instrument (Malvern). The hydrodynamic diameter and PDI were measured with a dispersion of 1 mg/mL NLCs in PBS while the zeta potential was measured with a dispersion of 1 mg/mL NLCs in 1 mM NaCl. Each assay was performed in three replications at 25°C.

### 2.1.5 Complexation of cNLCs with nucleic acid

In the complexation of cNLCs with model nucleic acid, all-star negative control siRNA (siMock) was carried out in PBS. The required volume for siMock was calculated according to the desired N/P ratios (ratio of positively-chargeable polymer amine (N = nitrogen) groups to negatively-charged nucleic acid phosphate (P) groups) at a constant concentration of the cNLCs nanocarrier (100 μg/mL). The cNLCs carrier and diluted siMock were gently homogenised by pipetting and kept for 10 min at room temperature before immediate use for downstream experiments.

### 2.1.6 Incubation with nanoparticles

For cell culture, 12, 24 and 96 cell culture microplates manufactured by Falcon^®^ or seahorse XFe96 were used. Cells were seeded at a concentration of 10^6^ cells/mL and cultured for 24 h. They were incubated for 24 h with nNLCs or cNLCs at a concentration ranging from 20 to 100 μg/mL. Cells were subsequently washed and stimulated with LPS (2 μg/mL) or IL-4 (20 ng/mL) for another 24 h. Finally, the impact of the two nanocarriers on BMDMs and BMDCs was assayed using various parameters, such as viability, phagocytosis, activation, cytokine secretion, nitric oxide (NO) production, reactive oxygen species (ROS) production and glycolysis or mitochondrial metabolism.

### 2.1.7 Toxicity assessment

Toxicity was measured by quantifying the cell viability using the CytoTox-ONE™ Homogeneous Membrane Integrity Assay kit (Promega, G7891) according to the manufacturer’s protocol. Briefly, the lysis solution (2 μl of lysis solution per 100 μl original volume) was used as a positive control for lactate dehydrogenase (LDH) release. A volume of 100 μL of CytoTox-ONE™ reagent was added to each well, before homogenisation on a shaker for 30 seconds and followed by incubation for another 10 min in the dark. After that, stop solution (50 μL) was added to each well, and the plate was placed on the shaker for another 10 seconds. Finally, their fluorescence was recorded at an excitation wavelength of 560 nm and an emission wavelength of 590 nm using a CLARIOstar^®^ microplate reader (BMG LABTECH).

### 2.1.8 Phagocytosis assay

Nanocarrier-exposed macrophages (BMDMs and J774.1A cells) and BMDCs were incubated at a ratio of 10 microspheres per cell for 6 h with 1.0-μm FluoSpheres^®^ carboxylate-modified microspheres (ThermoFisher, F8851) labelled with a red fluorescent dye (580 nm excitation and 605 nm emission). Cells were analysed by flow cytometry with an Accuri C6 instrument (Becton-Dickinson), and the analysis was performed by the FCS Express V5 software (De Novo Software).

### 2.1.9 Cell activation

Nanocarrier-exposed BMDCs and BMDMs were stimulated for 24 h using 2μg/mL LPS from *Escherichia coli*. Supernatants were collected for downstream cytokine immunoassay. After blocking the Fc receptor (BD Pharmingen, 553142) to reduce nonspecific binding, BMDCs and BMDMs were stained for CD11b (Ozyme, BLE101226) and CD11c (Ozyme, BLE117318) or CD11b (Ozyme, BLE101216) and F4/80 (Ozyme, BLE123152), respectively. To evaluate the cell activation, BMDCs and BMDMs were stained with anti-IAb (Ozyme, BLE116410) and CD86 (Ozyme, BLE105008) antibodies. In both cases, live cells were selected by negative 7-aminoactinomycin D (7AAD; BD Pharmingen, 559925) staining and analysed by flow cytometry using an LSR II instrument (Becton-Dickinson). The proportion of activated cells was quantified using FCS Express V5 software.

### 2.1.10 Cytokine immunoassays

Cytokine production was measured from cell culture supernatants with cytometric bead array (CBA; BD Pharmingen, 552364) using a mouse inflammation kit against IL-6, IL-12p70, MCP-1, TNFα, IL-10 and IFNγ. Results were acquired by flow cytometry using a BD LSR II instrument and analysed with FCAP Array Software v3.0 (BD Pharmingen, 652099).

### 2.1.11 NO and ROS Production

NO produced by BMDMs and BMDCs was determined by measuring nitrite concentration in cell culture media by Griess assay. Briefly, 50 μL of cell supernatant was transferred to a 96-well plate and incubated with an equal volume of sulphanilamide (Sigma, S9251) and N-alpha-naphthyl-ethylenediamine (Sigma, 222488) solutions, respectively, for 10 min each, protected from light. Optical density was measured at 540 nm using a CLARIOstar^®^ microplate reader, and sample nitrite concentration was determined using a standard curve. ROS production by BMDMs and BMDCs was determined by ROS-Glo™ H_2_O_2_ assay kit (Promega, G8821). The cells were cultured at 5 x 10^4^ cell/mL concentration in a 96-well plate, exposed to nanocarriers for 24 h and stimulated with 2 μg/mL of LPS. A volume of 20 μL of H_2_O_2_ substrate solution was added to each well before 6 h of ROS production measurement. ROS production measurement was performed by adding 100 μL of ROS-Glo™detection solution per well, before 20 min of incubation at 22°C followed by luminescence using a CLARIOstar^®^ microplate reader.

### 2.1.12 Metabolic flux analysis

For mature BMDCs (on day 10), 1.5 x 10^5^ cells per well were seeded into seahorse culture plate (Agilent, 102416-100) precoated with Cell-Tak (Corning, 354240) to enable BMDCs adherence, in complete culture media supplemented with GM-CSF (5 ng/mL) and FLT-3L (25 ng/mL). For mature BMDMs (on day 7), 0.8 x 10^5^ cells per well were seeded into seahorse culture plate as described in the previous study (Dey et al., 2021). A graphical representation of the experiment design is presented in Supplementary Figure 1.

### 2.1.13 Statistical analysis

Results are expressed as mean values ± standard deviation (SD). Statistical analysis was performed using GraphPad Prism version 8.4.2. Data were analysed by one-way ANOVA and Tukey’s multiple comparison test to analyse the difference between different groups. P-values below 0.05 were considered as significant and indicated as follows: *P ≤ 0.05, **P ≤ 0.01, ***P ≤ 0.001, and ****P ≤ 0. 0001 as compared with untreated cells (not exposed to NLCs).

## 3 Results

### 3.1.1 nNLCs and cNLCs do not induce cell toxicity and are efficiently internalised by APCs

We first investigated whether the exposure of nNLCs and cNLCs is toxic for APCs *in vitro,*using a macrophage cell line (J774.1A) or primary untransformed cells extracted from bone marrow: macrophages (BMDMs) and DCs (BMDCs). Cells were exposed to nNLCs or cNLCs with concentrations ranging from 0 to 250 μg/mL and measured toxicity. (Figure 1A). Among all the tested cells, BMDCs were most susceptible to both nNLCs and cNLCs exposure, and all the tested conditions exhibited more than 80% of cell viability. Therefore, for subsequent experiments, we chose 20 and 100 μg/mL as low and high standard doses, respectively, without adverse effects, that is, higher than 80% of cell viability after 24 h of incubation.

**Figure 1.**
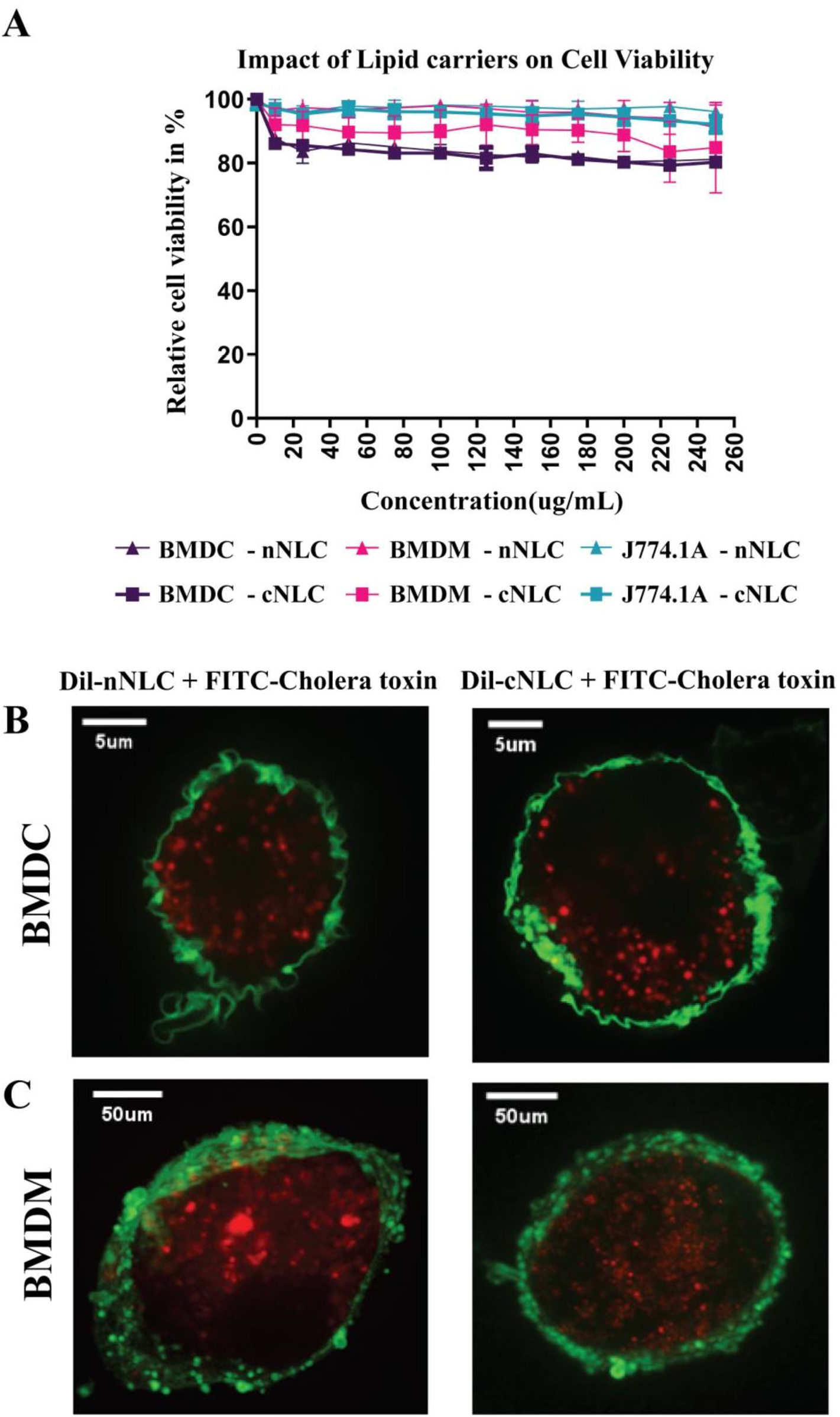
nNLCs and cNLCs do not induce cell toxicity and are efficiently internalised by APCs. **(A)** Cell viability (LDH release assay) of BMDCs, BMDMs and J774.1A cells was analysed after exposure to different concentrations of nNLCs and cNLCs nanocarriers for 24 h. Data are displayed as mean ± SD and normalised to the untreated cells (N = 3 independent experiments). Confocal microscopy analysis of nNLCs and cNLCs uptake in **(B)** BMDCs and **(C)** BMDMs. After APCs exposure to 100 μg/ml of nNLCs or cNLCs nanocarriers for 24 h, cell membranes were labelled with FITC-conjugated cholera toxin (green), and nNLCs and cNLCs are observed by excitation of Dil fluorescent dye (red). Images were acquired using a confocal spinning-disk microscope. The images displayed were representative of the majority of cells observed.

Next, we assayed the internalisation and cellular localisation of both nNLCs and cNLCs by two primary cell types: BMDCs and BMDMs that are more physiologically relevant than any immune cell lines. Both nanocarriers were internalised into the cytoplasm of BMDCs (Figure 1B) and BMDMs (Figure 1C) within a 24-h time frame. Therefore, from these first experiments, we can conclude that these two nanocarriers up to a 250-μg/mL concentration were not toxic, while they were both efficiently internalised by APCs.

### 3.1.2 nNLCs and cNLCs are internalised by APCs without affecting their phagocytic capacity

Accumulation of nanocarriers into phagocytic APCs opens the question of whether their functions could be altered, such as phagocytosis, which is one of the primary features of APCs. The phagocytic capacity of BMDCs or BMDMs was assessed by counting the number of engulfed microspheres per cell by flow cytometry. This parameter was not altered by either the neutral or the cationic nanocarrier supporting that the phagocytic capacity of both APCs was not modified by any type of nanocarrier (Figures 2A–2D). Moreover, we noticed that the phagocytic capacity of BMDMs was 20% higher than that of BMDCs (Figures 2B and 2D).

**Figure 2.**
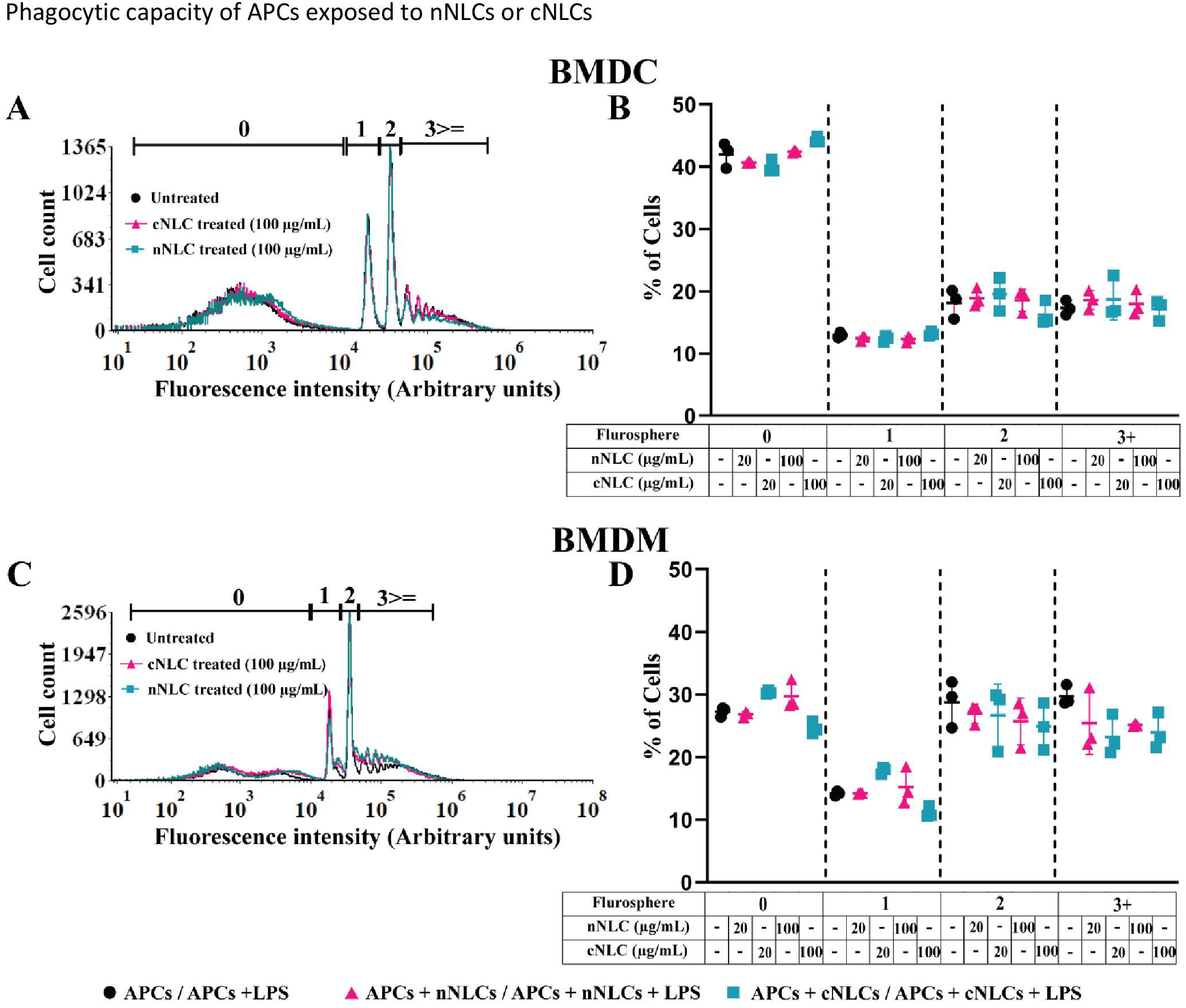
Phagocytic capacity of APCs exposed to nNLCs or cNLCs. BMDCs and BMDMs were exposed to nNLCs and cNLCs nanocarriers at 20 and 100 μg/mL for 24 h, then incubated with fluorescent microspheres for 6 h and subsequently analysed by flow cytometry. The repartition of the cells in the 1st, 2nd, 3rd and 4th peak corresponds to 0, 1, 2 and 3 or more beads internalisation, respectively. Overlaid histograms are shown in **(A)** for BMDCs and **(C)** for BMDMs. The proportion of cells in each peak was analysed for **(B)** BMDCs and **(D)** BMDMs. Data are displayed as mean ± SD (N = 3 independent experiments).

We also verified the impact of the nanocarriers on the phagocytic capacity of J774.1A cells, a well-characterised macrophage cell line for phagocytosis analysis (Luo et al., 2006). Similarly, we did not observe a significant change in phagocytic capacity between the nanocarrier treated cells or control cells. These results obtained with the J774.1A cell line were consistent with what we observed in the primary cells (Supplementary Figures 2A and 2B).

### 3.1.3 cNLCs but not nNLCs can increase LPS activation of BMDMs

BMDCs were identified by CD11b and CD11c expressions (Li et al., 2008) whereas BMDMs were marked by CD11b and F4/80 expressions (Zhang et al., 2008) (see the gating strategy in Supplementary Figure 3). Activation of BMDCs and BMDMs was evaluated by the frequency of CD86 and MHC-II double-positive cells. After LPS stimulation, the frequency of CD86^+^ and MHC-II^+^ in BMDCs increased from 27.83% to 75.9% (Figure 3A and Table 2) while no significative changes were observed in BMDMs (Figure 3B).

**Figure 3.**
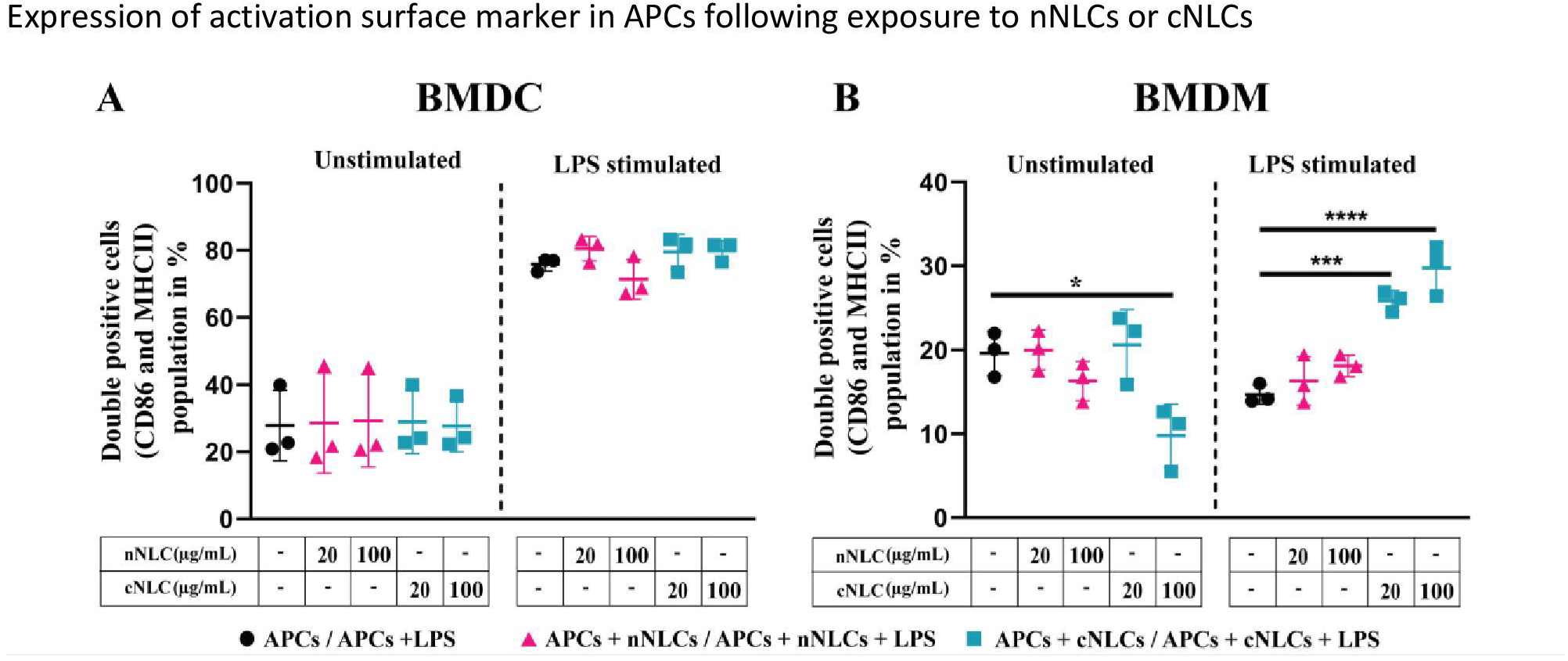
Expression of activation surface marker in APCs following exposure to nNLCs or cNLCs. BMDCs **(A)** and BMDMs **(B)** were exposed to nNLCs or cNLCs for 24 h, followed by LPS stimulation for an additional 24 h. Percentage of double-positive (CD86 and MHC-II) BMDCs and CD86 positive BMDMs were determined, with gating on CD11b and Cd11c positive cells for BMDCs and CD11b and F4/80 positive cells for BMDMs. Data are displayed as mean ± SD (N = 3 independent experiments), and the statistical significance between nanocarrier treated or untreated groups was performed by one-way ANOVA test using Tukey’s multiple comparisons test. *P ≤ 0.05; ***P ≤ 0.001; and ****P ≤ 0.0001.

**Table 2:** Percentage of activated APCs with or without NLCs treatment. Expression of activation surface marker of APCs. Expression of activation marker of BMDCs and BMDMs after exposure to nNLCs and cNLCs for 24 h, followed by LPS stimulation for another 24 h. Percentage of double-positive (CD86 and MHC-II) APCs were analysed. Prior to analyse, BMDCs were gated on CD11b^+^ and Cd11c^+^; BMDMs were gated on CD11b^+^ and F4/80^+^; and the data are presented in tabular form. Results are mean ± SD of 3 independent experiments.

|  | BMDCs |  | BMDMs |  |
| --- | --- | --- | --- | --- |
|  | Unstimulated | LPS-stimulated | Unstimulated | LPS-stimulated |
| Cells | $27.83 \pm 8.58$ | $75.9 \pm 1.62$ | $19.6 \pm 2.13$ | $14.69 \pm 0.93$ |
| Cells + nNLCs (20 ug/mL) | $28.61 \pm 12.22$ | $80.51 \pm 2.97$ | $19.98 \pm 1.92$ | $16.32 \pm 2.35$ |
| Cells + nNLCs (100 ug/mL) | $29.3 \pm 11.21$ | $71.38 \pm 4.85$ | $16.3 \pm 1.90$ | $18.1 \pm 1.05$ |
| Cells + cNLCs (20 ug/mL) | $28.97 \pm 7.79$ | $79.57 \pm 4.27$ | $20.61 \pm 3.39$ | $25.84 \pm 0.98$ |
| Cells + cNLCs (100 ug/mL) | $27.74 \pm 6.37$ | $79.91 \pm 2.39$ | $9.79 \pm 3.07$ | $29.76 \pm 2.45$ |

Exposure to increasing concentrations of nNLCs or cNLCs did not significantly alter LPS-induced double expression of CD86 and MHC-II in BMDCs. In the case of unstimulated BMDMs activation, CD86 and MHC-II double-positive cell percentage was not altered when exposed to nNLCs but decreased significantly when exposed to cNLCs at the highest dose from 19.6% to 9.79%. In the case of unactivated BMDMs, the percentage of CD86 positive cells remained unaltered when exposed to nNLCs (Table 2). Altogether, our data highlight that both nanocarriers do not activate BMDCs, but cNLCs slightly alter the activation of BMDMs. BMDCs, on exposure to both nanocarriers, maintained their capacity to respond to LPS activation. However, in the case of LPS-stimulated BMDMs, exposure to cNLCs significantly increased the percentage of activated BMDMs from 14.69% to 29.76%, while it remained the same with the nNLCs (Figure 3B and Table 2). This suggests that exposure to nanocarriers alone is not sufficient to activate both BMDCs and BMDMs. However, in LPS-stimulated BMDMs, exposure to cNLCs increased the frequency of CD86^+^ and MHC-II^+^ activated cells. Internalisation of both lipid nanocarriers, neutral and cationic ones, is not sufficient to activate both BMDCs and BMDMs, although exposure to cNLCs enhanced the ability of BMDMs to respond to LPS stimulation.

### 3.1.4 cNLCs and nNLCs can alter the production of signalling molecules by APCs

The capacity to produce different soluble factors, including signalling proteins such as cytokines or chemokines and other small molecular mediators such as NO and ROS, is a hallmark of APCs activation.

Having demonstrated that exposure to cNLCs could alter the activation of BMDMs in response to LPS, we wondered what would be the impact of both nanocarriers on cytokine secretion. We observed that both nanocarriers did not induce cytokine secretion in unstimulated BMDCs and BMDMs (Figures 4A–4D, left panel), except the highest dose of cNLCs but not nNLCs, which significantly increased the production of the MCP-1 chemokine in unstimulated BMDCs and to a lesser extend in unstimulated BMDMs (Figures 4E and 4F, left panel).

**Figure 4.**
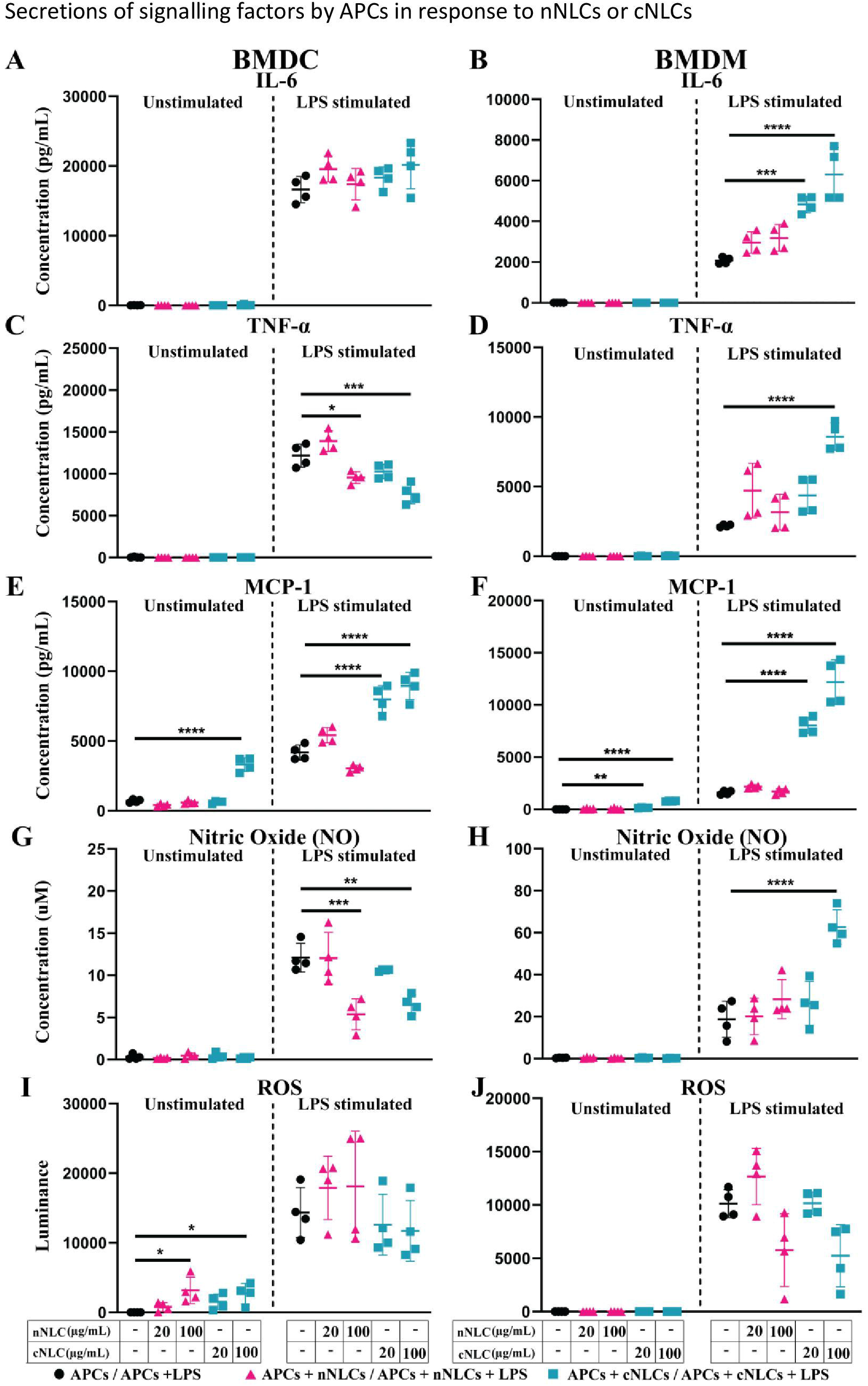
Secretions of signalling factors by APCs in response to nNLCs or cNLCs. Relative cytokine and chemokine concentration in the supernatant of BMDCs and BMDMs exposed to nNLCs or cNLCs and activated or not by LPS was determined by immunoassay. Secretion of the IL-6 cytokine in **(A)** BMDCs and **(B)** BMDMs; the TNFα cytokine in **(C)** BMDCs and **(D)** BMDMs and the chemokine MCP-1 in **(E)** BMDCs and **(F)** BMDMs. Relative NO concentration in the supernatant of BMDCs **(G)** and BMDMs **(H)** cells exposed to nNLCs or cNLCs and activated or not by LPS was determined by Griess assay. ROS production by BMDCs (I) and BMDMs (J) cells exposed to nNLCs or cNLCs and activated or not by LPS was determined by ROS-Glo™ H_2_O_2_ assay. Data are displayed as mean ± SD (N = 4 independent experiments), and the statistical significance between nanocarrier treated or untreated groups was performed by one-way ANOVA test using Tukey’s multiple comparisons test. *P ≤ 0.05; **P ≤ 0.01; ***P ≤ 0.001; and ****P ≤ 0.0001.

Upon LPS stimulation of APCs, nNLCs exposure did not alter IL-6 production by both BMDCs and BMDMs. However, exposure to cNLCs significantly increased IL-6 production by BMDMs (Figure 4B, right panel) but not by BMDCs (Figure 4A, right panel). In the case of BMDCs, both nNLCs and cNLCs decreased TNF-α production at 100 μg/mL (Figure 4C, right panel). For BMDMs, TNF-α production was only increased at 100 μg/mL of cNLCs but not for BMDCs (Figure 4D, right panel). We also observed that treatment with cNLCs but not nNLCs significantly increased MCP-1 production in both LPS-stimulated BMDCs and BMDMs (Figures 4E and 4F, right panel).

Two other important secretory molecules, NO and ROS productions were evaluated in the culture supernatant of APCs by Griess assay and H_2_O_2_ quantification, respectively. In absence of LPS stimulation, we did not observe a production of NO by BMDCs and BMDMs in response to both nanocarriers (Figures 4G and 4H, left panel) although ROS production was detected by BMDCs treated with 100 μg/mL of either nNLCs or cNLCs but not in BMDMs (Figures 4I and 4J, left panel). In LPS-stimulated conditions, both nNLCs and cNLCs at highest dose decreased NO production by BMDCs (Figure 4G, right panel), while the only cNLCs were responsible for increasing NO production in BMDMs (Figure 4H, right panel). After stimulation by LPS, both APCs produced increased quantities of ROS, but its production was not significantly altered by exposure to both nanocarriers (Figures 4I and 4J, right panel). These results indicate that BMDCs and BMDMs are differently affected by neutral or cationic nanocarriers regarding their capacity to produce NO and ROS and depending on activation stimuli.

Overall, nNLCs have only limited influence on the productions of signalling molecules, whereas cNLCs display significant effects, especially for inflammatory signals. The influence of cNLCs is clearly demonstrated in activated BMDMs by the increases of IL-6, TNF-α, MCP-1 secretions and NO production. Both nNLCs and cNLCs share most of their features such as their same size and composition; therefore, their major difference resides in their surface charge. This led us to hypothesise that this difference in the surface charge may be responsible for different effects driven by these two nanoparticles on APCs.

### 3.1.5 nNLCs and cNLCs have a significant impact on the mitochondrial metabolism of BMDMs but not on that of BMDCs

As cellular metabolism plays a key role in different functions of APCs, we sought to determine the effect of differentially charged LNCs on mitochondrial metabolism. For instance, pro-inflammatory stimuli by LPS are known to trigger a metabolic switch that would enhance glycolysis, whereas enhanced FAO and mitochondrial OXPHOS are hallmarks of IL-4-induced anti-inflammatory activity in immune cells.

Upon exposition to both nanocarriers, no alteration in the basal respiration, maximal respiration capacity, spare respiratory capacity, nonmitochondrial oxygen consumption and coupling efficiency (Supplementary Figures 4A, 4C, 4E, 5A, and 5C), proton leak or ATP production (Figures 5A and 5C) were found in unstimulated or stimulated BMDCs.

**Figure 5.**
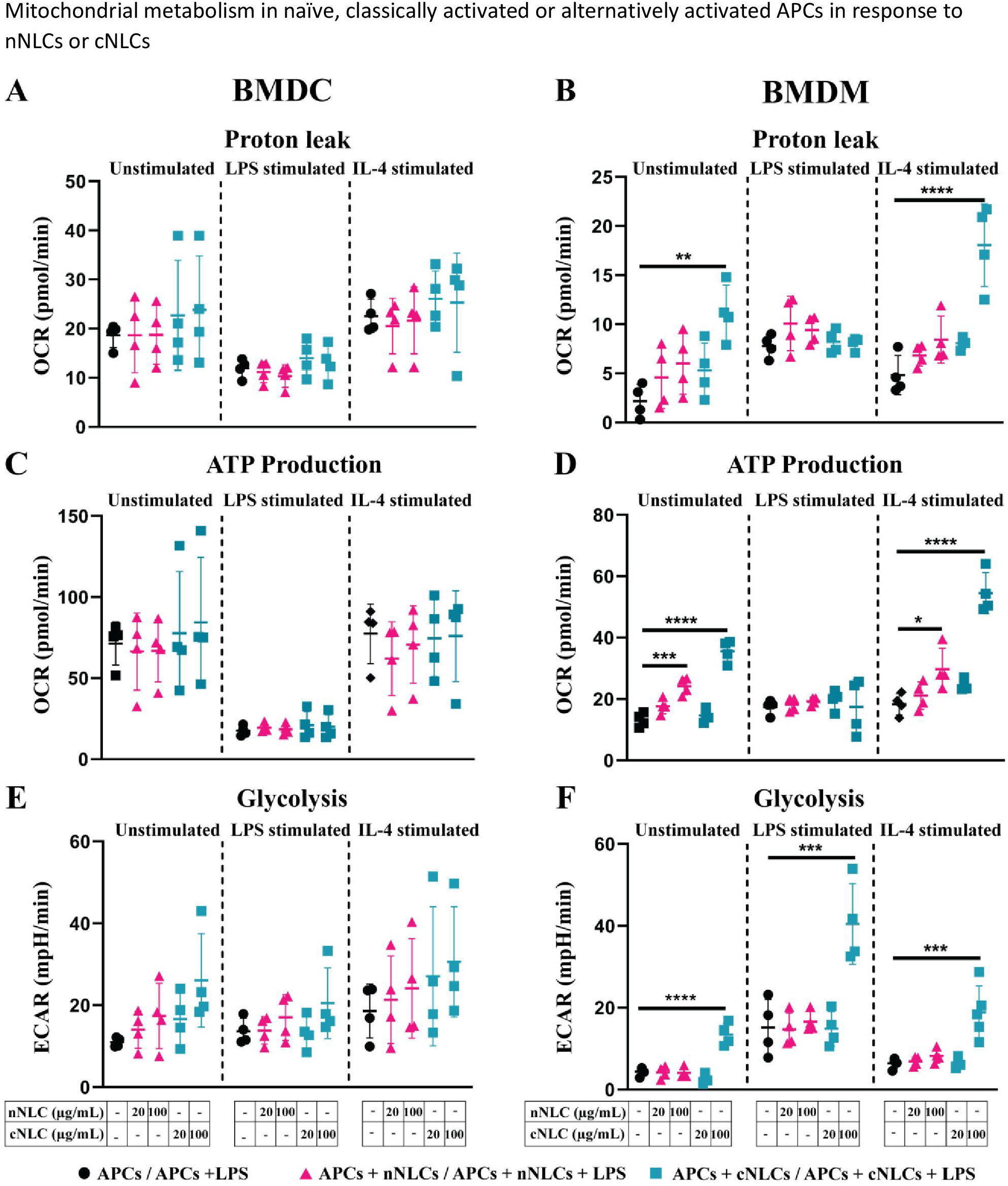
Mitochondrial metabolism in naïve, classically activated or alternatively activated APCs in response to nNLCs or cNLCs. **(A and B)** Proton leak, **(C and D)** ATP production and **(E and F)** glycolysis in BMDCs and BMDMs, respectively, were measured after exposure to cNLCs or nNLCs for 24 h and activated by LPS or IL-4 for another 24 h. Oxygen consumption rate (OCR) and ECAR were quantified using a seahorse XF analyser. Data were normalised by cell number based on cell count (Hoechst 33342 staining) and are displayed as mean ± SD (N = 4 independent experiments). The statistical significance between nanocarrier treated or untreated groups was performed by one-way ANOVA test using Tukey’s multiple comparisons test. *P ≤ 0.05; **P ≤ 0.01; ***P ≤ 0.001; and ****P ≤ 0.0001.

In BMDMs, exposure to both nanocarriers increased basal respiration and nonmitochondrial oxygen consumption of unstimulated cells at 100 μg/mL, as well as the nonmitochondrial oxygen consumption of LPS-stimulated cells treated with the nNLCs (Supplementary Figures 4B and 5B). Treatment with 100 μg/mL of cNLCs significantly increased the proton leak, Adenosine triphosphate (ATP) production, basal respiration, maximal respiration capacity, spare respiratory capacity and nonmitochondrial oxygen consumption (Figures 5B and 5D and Supplementary Figures 4B, 4D, 4F, and 5B) in unstimulated or IL-4-stimulated BMDMs whereas the nNLCs did only slightly increase basal respiration and nonmitochondrial oxygen consumption (Supplementary Figures 4A and 5A).

It is to be noted that both nanocarriers did not impair the coupling efficiency of unstimulated or stimulated BMDMs (Supplementary Figure 5B).

As a whole, our results demonstrate that the cNLCs have a more important effect on BMDMs’ metabolism compared with the nNLCs, while both nanocarriers have little effect on the metabolism of BMDCs.

### 3.1.6 nNLCs and cNLCs alter the glycolysis of BMDMs and not of BMDCs

Considering the alterations of the mitochondrial metabolism induced by the cNLCs and to a lesser extent the nNLCs, we sought to investigate their effects on the glycolytic profile of APCs as LPS-stimulated cells are mostly dependent on glycolysis. To evaluate the different glycolytic parameters of BMDCs and BMDMs, cells were first pretreated with different concentrations of both nanocarriers and then stimulated with LPS or IL-4 for 24 h. After stimulation, the extracellular acidification rate (ECAR) was measured using the glyco stress assay.

Unlike for BMDCs that did not show any alteration in glycolysis (Figure 5E) or glycolytic capacity (Supplementary Figure 6A), BMDMs’ glycolysis (Figure 5F) and glycolytic capacities (Supplementary Figure 6B) were increased in both unstimulated and stimulated conditions when exposed to 100 μg/mL of cNLCs. However, exposure to nNLCs did not induce any alteration in glycolysis or glycolytic capacity in BMDMs regardless of stimulating conditions (Figure 5F and Supplementary Figure 6B).

The combination of these results reveals that the cationic but not the nNLCs at the highest concentration alter the glycolytic profile in BMDMs. Conversely, both nanocarriers have no effect on glycolysis in BMDCs.

### 3.1.7 Reversing the surface charge with a nucleic acid cargo prevents adverse effects of cNLCs on APCs

As previous experiments have pointed out, at 100 μg/mL, cNLCs had a more dramatic effect on BMDMs’ physiology than nNLCs; we wondered whether the surface charge could explain the differences observed.

This led us to investigate whether we could reverse the phenotype observed on APCs by reversing the surface charge of the cNLCs with a nucleic acid cargo, here a negative control siRNA (siMock). We used different surface charges by fine-tuning the ratio of the positively charged amine groups of cNLCs nanocarriers (N = NH^3+^ group) relative to the negatively charged phosphate groups (P) from each phosphodiester bonds within the nucleic acid sequence, hence called N/P ratio. After complexation between siRNA and cNLCs nanocarriers, the zeta potential and hydrodynamic diameter of these nanocomplexes were measured. Naked cNLCs showed a zeta potential of 45.80 ± 3.8 mV in 1 mM NaCl while increasing amounts of the nucleic acid cargo and thus decreasing the N/P ratio lead to lower the zeta potential values down to –9.97 ± 0.94 mV, while naked nNLCs was measured at –16.50 ± 0.53 mV (Figure 6A). It is to be noted that the complexation of cNLCs with different quantities of siRNA did not significantly alter the size of the nanocomplexes (Figure 6B).

**Figure 6.**
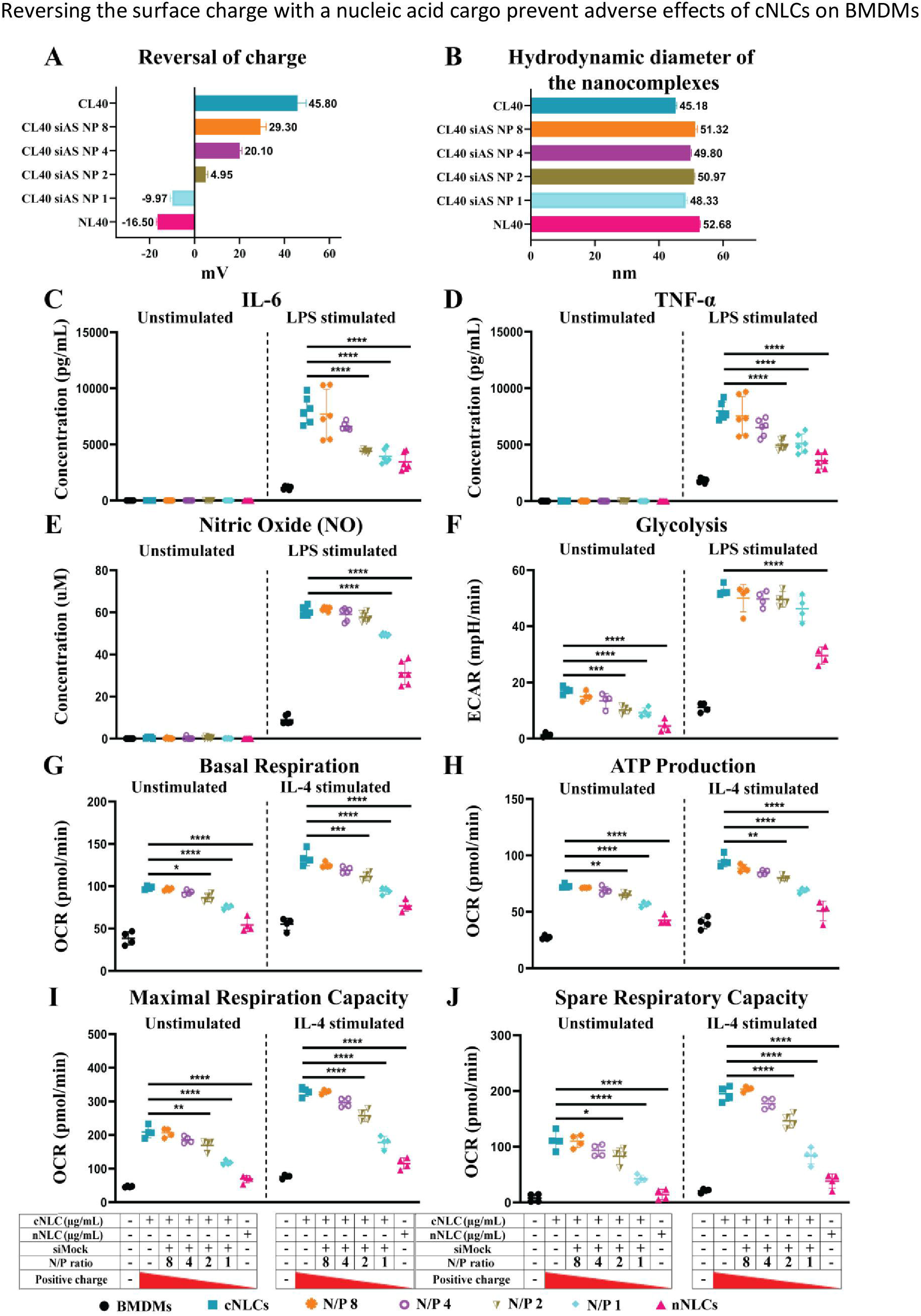
Reversing the surface charge with a nucleic acid cargo prevent adverse effects of cNLCs on APCs. **(A)** The zeta potential measurement of cNLCs complexes with siRNA at different N/P ratios was performed on a zetasizer instrument by ELS in 1 mM NaCl. **(B)** The hydrodynamic diameter of cNLCs complexes with siRNA at different N/P ratios was measured on a zetasizer instrument by DLS in PBS buffer. **(C)** IL-6 and **(D)** TNFα secretion was quantified from the supernatant of BMDMs exposed to 100 μg/mL of cNLCs complexes with siRNA at different N/P ratios and activated or not by LPS. **(E)** NO concentration in the supernatant of BMDMs exposed to 100 μg/mL of cNLCs complexes with siRNA at different N/P ratios and activated or not by LPS was determined by Griess assay. **(F)** Glycolysis in BMDMs exposed to 100 μg/mL of cNLCs complexes with siRNA at different N/P ratios and activated or not by LPS was determined by ECAR. **(G)** Basal respiration, **(H)** ATP production, **(I)** maximal respiration capacity and **(J)** spare respiratory capacity in BMDMs exposed to 100 μg/mL of cNLCs alone or complexes with siRNA at different N/P ratios and activated or not by IL-4 was determined by OCR. OCR and ECAR were quantified using a seahorse XF analyser. Data were normalised by cell number based on cell count (Hoechst 33342 staining) and are displayed as mean ± SD (N = 4 or 6 independent experiments). The statistical significance between nanocarrier treated or untreated groups was performed by one-way ANOVA test using Tukey’s multiple comparisons test. *P ≤ 0.05; **P ≤ 0.01; ***P ≤ 0.001; and ****P ≤ 0.0001.

Using different N/P ratios, we generated nanocarriers with different zeta potentials that we subsequently used to investigate their effects on BMDMs functions. An experimental design of metabolic flux analysis for reversal of nanocarrier surface charge is depicted in Supplementary Figure 7. BMDMs were exposed to 100 μg/mL of cNLCs nanocarrier, cNLCs-siRNA nanocomplexes at N/P 8 to N/P 1 or nNLCs nanocarrier. The culture supernatants were collected, and the secretion of pro-inflammatory cytokines (IL-6, TNFα) or chemokine (MCP-1) was quantified by immunoassay. IL-6 and TNFα productions by LPS-stimulated BMDMs were correlated to the zeta potential of the nanocarriers (Figure 6C and 6D), that is, the productions were maximum with cNLCs and decreased when cNLCs are complexed to siRNA reaching at N/P ratio 1 a similar level than the one obtained with nNLCs. The production of NO and MCP-1 by LPS-activated BMDMs also decreased with lower N/P ratios but to a lesser extent than for IL-6 and TNFα (Figure 6E and Supplementary Figure 8A).

To analyse the effect of the surface charge on glycolysis, we measured ECAR in BMDMs exposed to nanocomplexes at different N/P ratios and then stimulated or not with LPS. Both unstimulated and LPS-stimulated BMDMs showed a decrease in both glycolysis and glycolytic capacities with decreasing zeta potential and almost down to the same values as that of the nNLCs for the unstimulated cells (Figure 6F and Supplementary Figure 8B).

Next, we analysed the effect of the surface charge on the mitochondrial metabolism of BMDMs, by measuring the OCR in BMDMs exposed to nanocomplexes at different N/P ratios and then stimulated or not with IL-4. The exposure to differently charged nanocarriers showed a decrease in basal respiration, maximal respiration capacity, ATP production, spare respiratory capacity and proton leak correlated with a decrease in zeta potential in both unstimulated and IL-4-stimulated BMDMs (Figures 6G–6J and Supplementary Figure 8C). However, the effect of differently charged nanocarriers on both unstimulated and IL-4-stimulated BMDMs was not statistically significant for nonmitochondrial oxygen consumption and percentage of coupling efficiency (Supplementary Figures 8D and 8E).

Altogether, these results revealed that decreasing zeta potential, hence the surface charge of the cNLCs, was able to reverse their effect on the different cellular functions of primary BMDMs upon both pro- and anti-inflammatory stimulations. Moreover, using a range of N/P ratios representing the surface charge of the nanocarriers, we demonstrated that the alteration of the BMDMs physiology was proportional to the overall net surface charge of nucleic acid-loaded LNPs.

## 4 Discussion

Lipid-based nanocarriers are promising delivery systems for imaging (Navarro et al., 2012), gene therapy including nucleic acids delivery (Hibbitts et al., 2019) such as siRNA transfection (Bruniaux et al., 2014;Tezgel et al., 2018) or mRNA vaccine delivery (Zhang et al., 2019), drug delivery (Hinger et al., 2016), adjuvant delivery system (Bayon et al., 2018) and other biomedical applications.

Nanoparticles composed of cationic lipids have a strong capacity for binding and condensing nucleic acid by electrostatic interactions at the level of the phospholipid layer and deliver the payload across cellular membranes within the target cell cytoplasm (Elouahabi and Ruysschaert, 2005). However, when designing a lipid-based nanocarrier, the composition of the lipids defines the protein corona around the nanocarrier that is closely linked with the activation of the immune system leading to undesired side effects and biodistribution (Caracciolo et al., 2015;Moore et al., 2015). It is well known that different components of lipid-based carriers such as DOPE and DOTAP facilitate the formation of protein corona eventually causing undesired side effects (Caracciolo et al., 2013). One of the most efficient ways to reduce the nanocarrier-protein interaction and formation of protein corona is wrapping the nanocarrier with linear chains of PEG (Vonarbourg et al., 2006). PEGylation acts not only as an anti-opsonisation strategy but also as a thermodynamic shield that reduces nonspecific protein adsorption (Szleifer, 1997;Satulovsky et al., 2000). As our cNLCs contain DOPE and DOTAP, they were covered with 2 kDa PEG chains to limit the adsorption of proteins and direct interaction with plasma membrane as shown in a previous study(Wheeler et al., 1999), although preserving their capacity of the complexation with nucleic acids. However, it remains to assess the effects of cNLCs on different immune cells to precisely manage their future uses.

To understand the effect of differently charged NLCs, we opted for *ex vivo* experiments as an alternative to *in vivo* experiments, allowing for more regulated manipulation of cell functions and processes. Although cell lines have played a crucial role in scientific progress for decades, researchers are now increasingly skeptical when interpreting data generated from cell lines only. Factors such as misrepresented and contaminated cell lines have triggered a strong interest in primary cells (ATCC, 2010;Lorsch et al., 2014). In our study, to be closer to the physiological conditions, we conducted our experiments on BMDMs and BMDCs. Globally, in unstimulated BMDCs and BMDMs, NLCs had very few effects on the cellular production of soluble factors. Interestingly, after LPS stimulation, macrophages and DCs responded differently when treated with cNLCs and nNLCs. In the case of BMDMs, after LPS stimulation, cNLCs at high concentration provoked an enhanced immune response by increasing the production of different secretory pro-inflammatory molecules including IL-6, TNF-α, and MCP-1, while nNLCs did not. However, in the case of BMDCs, we observe a reduction in TNF-α secretion by nNLCs and cNLCs exposed LPS-stimulated. Under LPS stimulation, cNLC-exposed BMDCs and BMDMs increase their production of MCP-1. MCP-1 is one of the essential chemokines that governs the migration and infiltration of monocyte and macrophage (Deshmane et al., 2009). Elevations of MCP-1 production have been reported after the exposure of several nanomaterials such as gold NPs on BMDMs and BMDCs (Dey et al., 2021) or nickel NPs on mesothelial cells (Glista-Baker et al., 2012). Hence, MCP-1 may be considered as a sensitive indicator of NP exposure. MCP-1 is known to be associated with some inflammatory chronic diseases such as rheumatoid arthritis (Rantapää-Dahlqvist et al., 2007) or allergic asthma development (Ip et al., 2006). Therefore, it is important to consider the MCP-1 level when using cNLCs *in vivo* administration that might facilitate the emigration of immature myeloid cells at the site of exposure and promote inflammation.

To assess the influence of NLCs on the metabolism of BMDMs and BMDCs, we polarised these cells with either LPS or IL-4. While LPS-activated pro-inflammatory cells undergo a metabolic switch to enhanced glycolysis (Kelly and O’Neill, 2015;Van den Bossche et al., 2015), IL-4 induces alternatively activated cells towards an anti-inflammatory response, which would then rely mostly on FAO and mitochondrial OXPHOS (O’Neill and Pearce, 2016). As a result, altered metabolism is not only a characteristic of macrophage cell functions but also a prerequisite for a proper response to an immune stimulus. We demonstrated that both NLCs did not alter the basal mitochondrial respiration of BMDCs. However, in the case of BMDMs, basal respiration increased when exposed to the highest concentration used with both NLCs, indicating that the concentration of either neutral or cationic cargo must be finely determined. While no metabolic change was observed in BMDCs, they showed an increase of glycolysis and mitochondrial respiration specific of positive cNLCs. A previous study has shown a positive association between the glycolytic and the secretory activities in macrophages; however, the same was evaluated under LPS stimulation (Kelly and O’Neill, 2015). In unstimulated conditions with cNLCs exposure, we did not observe this coupling, probably because the cNLCs-induced increase of glycolysis is not high enough to drive secretory adaptations as observed in cNLCs-treated BMDMs under LPS stimulation. It is noteworthy that LPS-activated BMDMs rely on mitochondrial respiration. Based on these results obtained *in vitro*, we can assume that positive charge of cNLCs *in vivo* would not significantly affect the basal level of unstimulated DCs or macrophages secretory activity, hence preventing unintended immune responses (suppression or activation) and subsequent harmful outcomes (cancer or autoimmunity).

For our investigations, we used two NLCs with similar composition and size but solely differing by their zeta potentials. Therefore, the effects on the cellular functions of APCs observed only with cNLCs may be linked to their respective charge. This could be explained by three hypotheses: 1) the lipid composition of the NLCs (Caracciolo et al., 2015), 2) the net surface charge of NLCs (Fröhlich, 2012) and 3) the protein corona around NLCs (Henriksen-Lacey et al., 2011;Caracciolo et al., 2013). The pro-inflammatory effect of DOTAP-DOPE-based cNLCs has been previously documented in several studies explaining the interaction of cationic DOTAP with different immune cells. Here, we demonstrate that reversing the net charge of positively charged nanocarriers, by complexing with negatively charged siRNA, can reverse the effect of charged carriers on different cellular functions.

Therefore, we further studied the effect of the charge of the nanocarrier using BMDMs as a cellular model since they appeared to be the most affected cells by the exposure to cNLCs. By modifying the net surface charge of the cNLCs using negatively charged siRNA at different N/P ratios, we observed that the increase of the production of pro-inflammatory secretory molecules (IL-6, TNF-α, MCP-1 and NO) was proportional to the net surface charge of the lipid nanocarriers. In parallel, metabolic parameters, including basal respiration, maximal respiration capacity, ATP production, spare respiratory capacity and proton leak, were also modulated accordingly to the charge of the lipid nanocarriers. These results show that the effects of positively charged nanocarriers, such as cNLCs, can be reversed by the complexation of negatively charged ligands, such as RNA, proportionally to the net charge of the resulting nanocarrier. Different applications could then be developed with cNLCs associated with RNA, including RNAi therapeutics as well as mRNA delivery for vaccinal purposes, even in the context of immune disorders.

Several studies reported some effects of the charge of nanoparticles on cell behavior. For instance, N-Arginine-N-octyl chitosan is used to synthesise pH-sensitive charge-reversal lysosomolytic nanocarriers, which could reduce the potential toxicity of the nanocarrier as well as increase the drug delivery efficiency (Sun et al., 2017). Moreover, it has been shown that that charge-reversal nanocarriers enhanced gene delivery to the tumor site (Chen et al., 2016). Furthermore, researchers demonstrated that the use of chitosan and the pH-responsive charge-reversible polymer enhanced the siRNA delivery (Han et al., 2012). Here, our results highlight that fine-tuning of the surface charge of cationic NLCs with an oppositely charged biomaterial, for instance, nucleic acid, could prevent immunostimulation properties of the cationic carrier and has to be kept in mind for the future use of such carriers for therapeutic applications. Overall, using the same cationic lipid nanocarrier with tunable surface charge, we propose that positive charge is one of the major factors responsible for the alteration of the immune response.

## 5 Conclusion

In conclusion, both BMDCs and BMDMs responded differently when exposed to the cationic or neutral variation of the same lipid nanocarriers. Therefore, it is highly relevant to include both cell types in the case of immunotoxicity analysis. We demonstrated that both nanocarriers, at low concentration, did not significantly alter several functions of both APCs. However, the cationic nanocarrier, at the highest concentration, induced alterations of some functions of APCs. We demonstrated that this effect on APCs was dependent on the net positive charge surface charge of the lipid carrier that could be offset by loading nucleic acid cargo that mediated reversal of the charge. Finally, we propose that tuning the nucleic acid load, hence, the surface charge of NLCs is critical to their use for therapy and prevent the alteration of immune cell response to stimuli.

## 7 Author Contributions

AKD, AN, FC, CF, FPN and PNM wrote the manuscript. AN, DJ, ME and FPN synthesized the nanoparticles and performed their physico-chemical characterization. AKD, CF and PNM designed and performed cell experiments. FC, EJM, FPN and PNM analysed the data and reviewed the study.

## 9 Acknowledgments

The authors acknowledge the staff of the animal facility of IAB, C. Charrat for technical support, M. Pezet for confocal and flow cytometry analysis, S. Blanchet for her expertise in SeaHorse analysis, Z. Macek-Jilkova for stimulating discussions. This publication reflects only the author’s view and the Commission is not responsible for any use that may be made of the information it contains.

---

Supplementary Figure 1 | Experimental design of metabolic flux analysis

Mature BMDCs and BMDMs were seeded on a Seahorse culture plate. One hour after plating, cells were treated with the different nanocarriers. After 24 h of culture, cells were washed and when indicated, stimulated with LPS or IL-4 for 24 h. The metabolic analysis was performed using a Seahorse bio-analyser using the Mito Stress and Glyco Stress assay protocol, with the corresponding chemical inhibitors.

Supplementary Figure 2 | Phagocytosis capacity of macrophage cell line J774.1A

J774.1A cells were exposed to nNLCs and cNLCs nanocarriers at 100 μg/mL for 24 h, then incubated with fluorescent microspheres for 6 h, and subsequently analysed by flow cytometry. The repartition of the cells in the 1st, 2nd 3rd and 4th peak corresponds to 0, 1, 2 and 3 or more beads internalization, respectively. Overlaid histograms are shown in **(A)** The proportion of cells in each peak was analysed **(B)**. Data is displayed as mean ± SD (N = 3 independent experiments).

Supplementary Figure 3 | Expression of activation surface marker in APCs

The expression of activation marker for BMDCs and BMDMs was quantified by flow cytometry after exposure to nNLCs or cNLCs for 24 h, followed by LPS stimulation for additional 24 h when indicated. The percentage of double positive (CD86 and MHC-II) BMDCs and BMDMs were gated on CD11b and Cd11c positive cells for BMDCs and CD11b and F4/80 positive cells for BMDMs and contour graph was displayed. The results are representative one of the three independent experiment.

Supplementary Figure 4 | Basal respiration, maximal respiration capacity, and spare respiratory capacity of naïve, classically activated or alternatively activated APCs in response to nNLCs or cNLCs.

**(A, B)** Basal respiration, **(C, D)** Maximal respiration capacity, **(E, F)** Spare respiratory capacity of BMDCs and BMDMs respectively were measured after exposure to cNLCs or nNLCs for 24 h and activated by LPS or IL-4 for another 24 h. Oxygen consumption rate (OCR) was quantified using a Seahorse XF analyser. Data was normalized by cell number based on cell count (Hoechst 33342 staining) and is displayed as mean ± SD (N = 4 independent experiments). Statistical significance between nanocarrier treated or untreated groups was performed by one-way ANOVA test using Tukey’s multiple comparisons test. *P ≤ 0.05; **P ≤ 0.01; ***P ≤ 0.001; ****P ≤ 0.0001.

Supplementary Figure 5 | Non-mitochondrial oxygen consumption and percentage of coupling efficiency of naïve, classically activated or alternatively activated APCs in response to nNLCs or cNLCs.

**(A, B)** Non-mitochondrial oxygen consumption, **(C, D)** percentage of coupling efficiency of BMDCs and BMDMs, respectively, were measured after exposure to cNLCs or nNLCs for 24 h and activated by LPS or IL-4 for another 24 h. Oxygen consumption rate (OCR) was quantified using a Seahorse XF analyser. Data was normalized by cell number based on cell count (Hoechst 33342 staining) and is displayed as mean ± SD (N = 4 independent experiments). Statistical significance between nanocarrier treated or untreated groups was performed by one-way ANOVA test using Tukey’s multiple comparisons test. *P ≤ 0.05; **P ≤ 0.01; ***P ≤ 0.001; ****P ≤ 0.0001.

Supplementary Figure 6 | Glycolytic capacity of naïve, naïve, classically activated or alternatively activated APCs in response to nNLCs or cNLCs.

Glycolytic capacity **(A)** in BMDCs and **(B)** in BMDMs were evaluated after exposure to cNLCs or nNLCs for 24 h and activated by LPS or IL-4 for another 24 h. Extracellular acidification rate (ECAR) was quantified using a Seahorse XF analyser. Data was normalized by cell number based on cell count (Hoechst 33342 staining) and is displayed as mean ± SD (N = 4 independent experiments). Statistical significance between nanocarrier treated or untreated groups was performed by one-way ANOVA test using Tukey’s multiple comparisons test. **P ≤ 0.01; ***P ≤ 0.001; ****P ≤ 0.0001.

Supplementary Figure 7 | Experimental design of metabolic flux analysis for reversal of nanocarrier surface charge.

Mature BMDCs and BMDMs were seeded in Seahorse culture plate. One hour after plating, cells were treated with the different nanocarriers, and when indicated with nanocarriers/siRNA nanocomplexes at the corresponding N/P ratios. After 24 h of culture, cells were washed and when indicated stimulated with LPS or IL-4 for 24 h. The metabolic analysis was performed using a Seahorse bio-analyser using the Mito Stress and Glyco Stress assay protocol, with the corresponding chemical inhibitors.

Supplementary Figure 8 | Effect of the net surface charge of cNLCs on different cellular functions and metabolism of BMDMs

**(A)** MCP-1 production was quantified from the supernatant of BMDMs exposed to 100 μg/mL of cNLCs complexes with siRNA at different N/P ratios and activated or not by LPS. **(B)** Glycolytic capacity in BMDMs exposed to 100 μg/mL of cNLCs complexes with siRNA at different N/P ratios and activated or not by LPS was determined by extracellular acidification rate (ECAR). **(C)** Proton leak **(D)** non-mitochondrial oxygen consumption, **(E)** percentage of coupling efficiency in BMDMs exposed to 100 μg/mL of cNLCs complexes with siRNA at different N/P ratios and activated or not by IL-4 was determined by Oxygen consumption rate (OCR). OCR and ECAR were quantified using a Seahorse XF analyser. Data was normalized by cell number based on cell count (Hoechst 33342 staining) and is displayed as mean ± SD (N = 3 independent experiments). Statistical significance between nanocarrier treated or untreated groups was performed by one-way ANOVA test using Tukey’s multiple comparisons test. *P ≤ 0.05; **P ≤ 0.01; ***P ≤ 0.001; ****P ≤ 0.0001.

## 12 Data Availability Statement

Due to confidentiality agreements, supporting data can only be made available to bona fide researchers, subject to a nondisclosure agreement.

